# Hoisted with his own petard: how sex-ratio meiotic drive in *Drosophila affinis* creates resistance alleles that limit its spread

**DOI:** 10.1101/2022.02.14.480432

**Authors:** Wen-Juan Ma, Emma M. Knoles, Kistie B. Patch, Murtaza M. Shoaib, Robert L. Unckless

## Abstract

Meiotic drivers are selfish genetic elements that tinker with gametogenesis to bias their own transmission into the next generation of offspring. Such tinkering can have significant consequences on gametogenesis and end up hampering the spread of the driver. In *Drosophila affinis*, sex-ratio meiotic drive is caused by an X-linked complex that, when in males with a susceptible Y chromosome, results in broods that are typically more than 95% female. Interestingly, *D. affinis* males lacking a Y chromosome (XO) are fertile and males with the meiotic drive X and no Y produce only sons - effectively reversing the sex-ratio effect. Here, we show that meiotic drive dramatically increases the rate of nondisjunction of the Y chromosome (at least 750X), meaning that the driver is creating resistant alleles through the process of driving. We then model how the O might influence the spread, dynamics and equilibrium of the sex-ratio X chromosome. We find that the O can prevent the spread or reduce the equilibrium frequency of the sex-ratio X chromosome and it can even lead to oscillations in frequency. Finally, with reasonable parameters, the O is unlikely to lead to the loss of the Y chromosome, but we discuss how it might lead to sex-chromosome turnover indirectly.

## 1 INTRODUCTION

Selfish genetic elements enhance their own transmission to the next generation, which can be neutral or detrimental to host fitness (Werren et al., 1988; Werren, 2011). As a consequence, genetic conflict occurs between the selfish genetic elements and other genomic regions. Such selfish genetic elements, including transposable elements, homing endonucleases, meiotic drivers and heritable microorganisms, are ubiquitous in eukaryotes with new cases reported regularly (Hurst and Werren, 2001; Burt and Trivers, 2006). This conflict can have profound evolutionary consequences ranging from speciation to chromosome evolution to extinction (Burt and Trivers, 2006; Lindholm and Price, 2016). In response to such genomic conflict, resistance alleles may evolve to counterbalance the selfish genetic elements and associated fitness costs (Hall, 2004; Courret et al., 2019; Price et al., 2020). However,cases where the selfish genetic elements actually create resistance alleles themselves are rare (but see Bravo Nũnez *et al*. 2020).

Meiotic drivers are selfish genetic elements that promote their own representation in gametes, resulting in highly biased transmission (Lindholm et al., 2016; Zanders and Unckless, 2019). Sex-ratio (SR) meiotic drive occurs when a drive element on the X (or much more rarely Y) chromosome manipulates gametogenesis to prevent the maturation of Y-bearing sperm (or X-bearing) in males, resulting in the production of predominantly female (or male) offspring (Jaenike, 2001) (Figure 1, parameters described in Table 1). Most *Drosophila* species requires a Y for fertility - XO males (lacking a Y) are often sterile (Pomiankowski et al., 2004). There are a handful of exceptions to XO sterility in *Drosophila*, and *Drosophila affinis* is one of them - *D. affinis* XO males are fertile. However, XO males tend to have a significant fitness cost, with the severity of the cost dependent on the genetic background (Voelker and Kojima, 1972; Unckless et al., 2015). In *D. affinis*, SR meiotic drive occurs in various populations across the Eastern United States (Voelker, 1972; Unckless et al., 2015), with an estimated frequency of 0.02-0.1 (personal observations). Y-chromosome linked resistance has evolved to counterbalance the extreme female-biased sex ratio induced by meiotic drive (Voelker, 1972; Unckless et al., 2015). What makes *D. affinis* unique is that males with sex-ratio X chromosome and O (X^SR^O males) sire only sons, effectively reversing drive (Voelker (1972), Figure 1).

**FIGURE 1.**
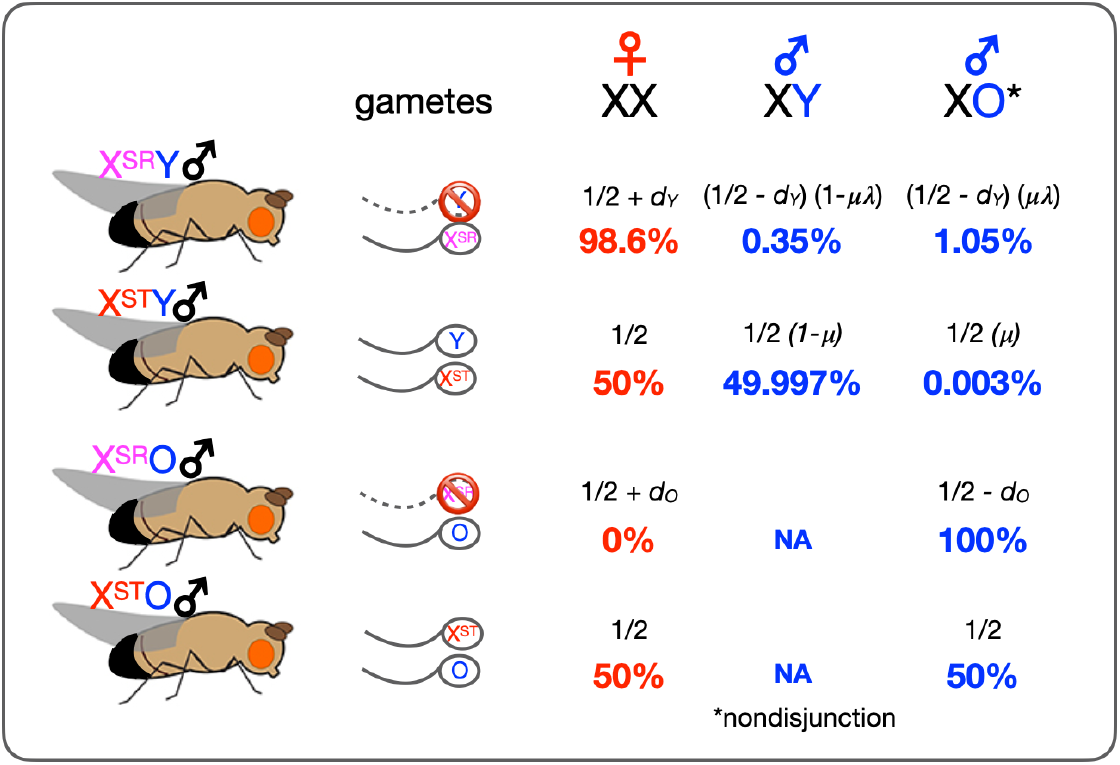
An overview of SR meiotic drive and nondisjunction rate in males in *D. affinis*. ST: Standard (non-driving) strain, SR: sex-ratio strain, O: missing Y chromosome. Solid line indicates gamete produced and dotted line refers to nonviable gamete. Red numbers refer to female offspring percentage, blue numbers refer to male offspring (estimates from empirical data). Formulas refer to model estimates for offspring produced. *d*_*Y*_, *λ, µ*, are model parameters described in Table 1.

**TABLE 1.**
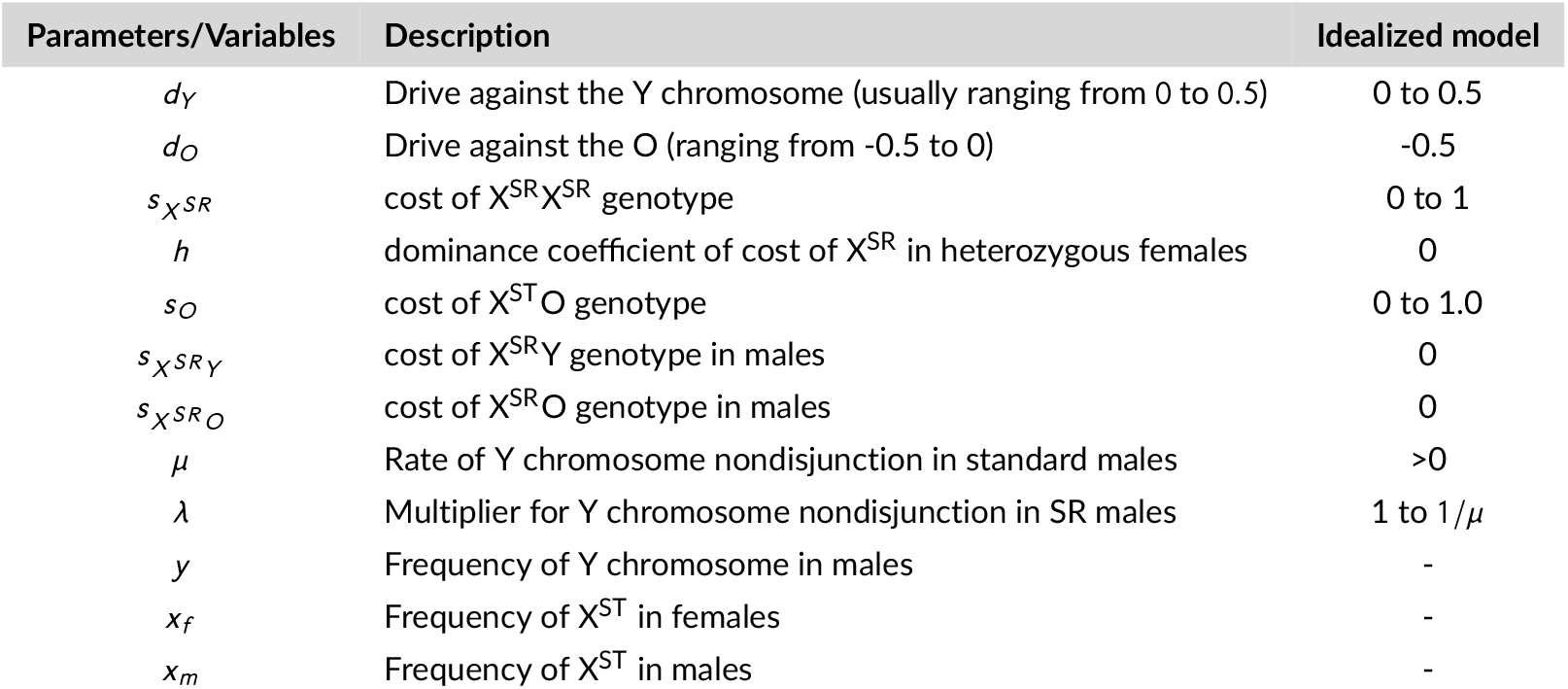
Model parameters and variables

XO males arise through nondisjunction which occurs when homologous chromosomes fail to segregate properly during meiosis, resulting in the gametes with an improper chromosome complement (Day and Taylor, 1998). Sex chromosome nondisjunction can dramatically affect the sex determination or sexual development in organisms, and *Drosophila* is one of the most intensely studied systems on sex chromosome nondisjunction (Bridges, 1913; Kelsall, 1961; Tokunaga, 1970; Koehler et al., 1996; Xiang and Hawley, 2006). Though there are essential fertility factors on the Y in most *Drosophila* species, sex is determined by the ratio of autosomes to the X (Pomiankowski et al., 2004). Thus in *Drosophila* species, meiotic nondisjunction in both sexes could result in viable XXY daughters and XO sons (Belote and Lucchesi, 1980; Cheng and Disteche, 2006). However, flies with a single Y chromosome (YO) or 3 X chromosomes are inviable (Cheng and Disteche, 2006). In both *D. pseudoobscura* and *D. neotestacea* (two well-studied systems with SR meiotic drive), the sons produced by SR males are invariably XO and sterile (Henahan and Cobbs, 1983; James and Jaenike, 1990; Dyer, 2012).

Selfish genetic elements were early inspiration for population manipulation strategies (Hastings, 1994; Ribeiro and Kidwell, 1994). Recently, synthetic homing gene drives have garnered attention as a possible means of manipulating populations of organisms including mosquitoes (Burt, 2003; Deredec et al., 2008; Hammond et al., 2016; Champer et al., 2016). These homing gene drive systems function by targeting a specific sequence motif that is then cut and repaired either through homology directed repair (insertion of the gene drive machinery) or nonhomologous end joining which often leads to the creation of a resistance allele (Champer et al., 2016). These resistance alleles arise because nonhomologous end joining results in mutations in the target sequence rendering the target immune to future homing. While more recent strategies aim to mitigate this type of resistance, it remains a basic problem with homing-style gene drives.

These synthetic gene drive systems can and should be informed by natural drive systems (including SR meiotic drive) because the natural systems are already natural experiments (Price et al., 2020). Likewise, synthetic gene drives can inform our understanding of natural systems since we can watch them spread through (experimental or even natural) populations from introduction to fixation, loss or other outcomes. The drive system in *D. affinis* provides an interesting corollary to what happens with homing gene drives. In both systems, resistance alleles occur at some baseline rate: standard nucleotide mutations or indels for homing gene drives and nondisjunction of the Y chromosome in *D. affinis*. However, the drive machinery creates these resistance alleles at rates orders of magnitude higher due to the mechanisms of drive. Understanding the evolution and mechanisms of resistance is crucial for better engineering effective gene drive systems (Price et al., 2020).

Meiotic drive has been theorized - and in some cases shown empirically - to have significant ecological and evolutionary consequences for populations (Lyttle, 1991; Jaenike, 1999, 2001; Reinhardt et al., 2014; Unckless et al., 2015). One such consequence is the turnover of sex chromosomes similar to what was theorized by Van Doorn and Kirkpatrick (2007) for sexual conflict in general. Given the fact that XO males are fertile in *D. affinis*, this raises the possibility that the O could lead to extinction of the Y chromosome, extinction of the X^SR^ chromosome (X chromosome with sex-ratio meiotic drive) or both. We first empirically measured rates of Y chromosome nondisjunction in *D. affinis* in standard (ST) and SR males and the frequency of XO males in the wild. This provided an estimate of how nondisjunction rates are influenced by SR meiotic drive and allowed us to use population genetic modeling to address several questions. First, we investigated whether the segregation of the O could prevent invasion of X^SR^ chromosome. Second, we examined the equilibrium frequencies of the O and X^SR^ under a range of parameters, and specifically compared the frequency of X^SR^ in the presence and absence of the O in a population. We found that often the O reduced X^SR^ frequency considerably, especially when the cost of O was low. Finally, we considered under what circumstances the O and/or X^SR^ might lead to the complete loss of the Y chromosome. Our models do not include the Y chromosome resistance and the effects on the dynamics of X^SR^, because this has been extensively modeled in Hall’s model (2004). In addition, adding resistant Y chromosomes dramatically increases the model complexity, for instance, including two X chromosomes, and 3 Y chromosome status. Rather, our models expand Hall’s model (2004) to include several additional male fitness costs and consider the specific role of the O. The complexity of the full model precluded a closed form solution for equilibrium, so we considered an idealized model where several parameter values were fixed. This allowed us to find analytical solutions and, in some cases, train our intuition. We then used deterministic simulations to explore parameter space more broadly.

## 2 MATERIALS AND METHODS

### 2.1 Fly stocks and husbandry

We utilized three lab stocks of *D. affinis* in these experiments. One standard (non-driving) stock is referred to as *darkeye* strain because it carries an X-linked recessive dark eye color mutation. It was collected in Rochester, NY in 2012 and inbred for 10 generations. The second standard stock was obtained from the predecessor of the National *Drosophila* Species Stock Center (14012-0141.02), and is referred to as the *genome* strain (in preparation). It was collected in Halsey National Forest in Nebraska in 1958. The sex-ratio stock (SR18) was collected in Rochester, NY in 2018 and backcrossed to the *darkeye* strain for at least 10 generations (see Unckless et al. (2015)). Thus, both the SR and *darkeye* stocks should carry the *darkeye* autosomes and Y chromosome. Flies were maintained on malt agar food (80g malt extract, 60g semolina flour, 20g nutritional yeast and 10g agar in 1.05l of water supplemented with both 14ml 10% Tegosept solution and 5ml Propionic acid) in clear, polystyrene vials (95mm by 25.12mm) and dry yeast powder was sprinkled on the medium surface to stimulate female egg laying. Cotton rolls were added when just prior to pupation and serve as a substrate for pupation. Flies were maintained at constant temperature at 21°C in a 12 hour light/dark cycle.

Wild males were collected at Rees’ Fruit Farm (39.091 latitude, -95.594 longitude) near Topeka, KS in August 2021 using bottle traps baited with banana mash, using similar collecting method described in (Gleason et al., 2019). Traps were hung in the branches of two apricot trees or in a raspberry hoophouse for 3-4 days, and all live *Drosophila* flies were transferred to polypropylene *Drosophila* bottles (177ml) with standard food via aspiration. We then immediately separated *D*.*affinis* from other species under stereo binocular microscope (Stereo microscope z850, Chesterfield, MO, USA) in the lab.

### 2.2 Rates of nondisjunction in standard and sex-ratio males in the lab and the frequency of the O in the wild population

To investigate rates of Y chromosome nondisjunction in sex-ratio males, 1-3 virgin females (1-4 days old) from the *darkeye* strain were crossed with 1 SR18 male (7-10 days old) for each replicate. Similarly, for Y chromosome nondisjunction rates in standard (the *darkeye* strain) males, 1-3 virgin females (1-4 days old) from the *genome* strain were crossed with 1 *darkeye* (ST) (7-10 days old) for each replicate. The crossing scheme was set up to distinguish Y chromosome nondisjunction from potential rare X chromosome nondisjunction, taking advantage of the eye color polymorphism between the two strains. To mitigate potential environmental factors, the two crosses were performed at the same time, with strict controlled culture condition (at 21°C in a 12 hour light/dark cycle). We also measured the frequency of XO in the wild by PCR screening (see below) the males collected from the Rees’ Fruit Farm.

### 2.3 Genomic DNA extraction and XO male frequency screening

We isolated genomic DNA of each individual male using a modified Puregene Gentra Tissue kit (Qiagen, Hilden, Germany). Briefly, each individual fly was placed in a well of a 96-well DNA extraction plate, with one 2.5mm glass beads and 100µl cell lysis solution before homogenizing in a bead beater (Mini Beadbeater™, Biospec Products INC., Bartlesville, OK, USA). The rest of the protocol followed the standard Puregene Gentra Tissue kit manual.

Primers (Table S.1) to amplify a Y-specific DNA fragment were developed using transcripts from male testes blasted against a female genome assembly (in preparation). PCRs were performed in a total volume of 10µl (1µl DNA sample, 3µl Milli-Q water, 5µl GOTag Green Master Mix buffer (Promega, note with 1.5mM MgCl2 for 1x reaction), 0.5µl forward primer and of reverse primer each). PCRs were conducted on Bio-Rad T100 Thermal Cycler (Bio-Rad Laboratories, Hercules, CA, USA) using the following profile: 2 min of Taq polymerase activation at 95°C, followed by 35 cycles including denaturation at 95°C for 20 sec, annealing at 56°C for 30 sec and elongation at 72°C for 45 sec, followed by a final elongation of 5 min at 72°C. Afterwards, 3-5µl of PCR product was mixed with 3µl of loading dye and loaded to run a 1.5% agarose gel. We repeated negative PCRs 1 to 4 times for Y-linked gene PCRs to confirm that these were indeed XO males.

For individual male samples, we also performed PCR on the COI gene (primers in Table S.1) to assure DNA quality. Only those male samples negative for the Y-specific marker but positive for COI were considered as XO males. The PCR master mix for COI was the same to those of Y-specific marker. The PCR protocol was identical except that the annealing temperature was 55°C.

### 2.4 Model

Our full model is described in the Appendix, but is an expansion of what was developed by Hall (2004). While our recursions use the frequency of the standard X^ST^ and Y chromosomes, our plots consider the frequency of the sex ratio X^SR^ chromosome and O. The substantive differences between the Hall model and our model are a) that X^SR^O males can produce anywhere from all *O* -bearing sperm (*d*_*O*_ = −0.5) to all *X* ^*SR*^ -bearing sperm (*d*_*O*_ = 0.5), though we focus on the case where *d*_*O*_ = −0.5 (observed in *D. affinis* (Voelker, 1972; Unckless et al., 2015)), b) the various male genotypes carry costs 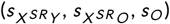, and c) that nondisjunction converts Y to O at rate *µ* in XY males and *µ* ∗ *λ* in X^SR^Y males. So, *λ* is a multiplier that can range from 1 (the same rate for SR and standard males) to 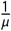 (the rate of nondisjunction in SR males is one). All parameters are described in Table 1.

We were unable to derive analytical solutions for the full model, so we developed an idealized model where we fix the values of several parameters, which allowed for some analytical progress. We refer to this throughout as the “Idealized” model, but use deterministic simulation to explore the full model. Analytical solutions were derived using Mathematica (Version 12, Wolfram|Alpha, Champaign, Illinois, USA).

### 2.5 Simulations

We use deterministic forward simulations to check our analytical solutions and to explore the full parameter space. All simulations were coded in R (R Core Team, 2018) and code is available through Github (https://github.com/unckless/SROdynamics) Briefly, simulations were initiated with *x*_*f*_ = 0.999, *x*_*m*_ = 0.999, and 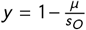 (reflecting mutation-selection balance of the O) with an effectively infinite population size. Each generation, new frequencies were calculated and recorded using the recursions in Appendix Equations (A.1)-(A.3). This continued for 20000 generations, after which we recorded a) the frequencies of the standard X and X^SR^ chromosomes in females and males, b) the frequencies of Y chromosome and O in males, c) the stability of the frequencies (binary based on whether the variance over the final 2000 generations was less than 0.001) and d) whether or not the X^SR^ and/or O invaded.

### 2.6 Data analysis

Data analysis and figure generation were performed in R (R Core Team, 2018), utilizing several packages including ggplot2 (Wickham, 2011), cowplot (Wilke et al., 2019), metR (Campitelli, 2021), dplyr (Wickham et al., 2021), and forcats (Wickham, 2019). Nondisjunction rates among standard and SR meiotic drive males were compared using Fisher’s exact test, and XO male frequency in the wild males was compared to expected frequency by mutationselection balance with Chi-Square test.

## 3 RESULTS

### 3.1 Empirical evidence that SR meiotic drive leads to nondisjunction of the Y in *D. affinis*

#### 3.1.1 The rate of nondisjunction in standard and sex-ratio males in the lab

In other species, sons of SR fathers are invariably XO because they lose the Y during gametogenesis (Henahan and Cobbs, 1983; James and Jaenike, 1990; Dyer, 2012). We determined the rate of Y chromosome nondisjunction in SR and ST males of *D. affinis* using a PCR assay. In total, 51 SR fathers produced broods. Note that one clutch had only 1 daughter and 1 son, but this son is an XO male (i.e. clutch sB26, Table S.2). Among the 51 broods, approximately 1.4% (69 out of 4786) of the offspring were male (ranging from 0 to 23.8% per brood). Among the 51 broods, 26 included at least one son, with on average 2.30 (SE=0.05, Table S.2). We performed PCR for a Y-specific marker on 56 of these sons, and 75% (42 out of 56, with average proportion of 77.5% across individual fathers (SE=O.O75)) were XO (repeated negative for Y-specific marker but positive for COI marker, Table 2). These XO males were all *darkeye* males, consistent with them losing their Y chromosome during gametogenesis. Nondisjunction in mothers could lead to XO sons if the father contributes the X, but with our crossing scheme, those sons would have wild type eyes.

**TABLE 2.** Rates of nondisjunction and XO male frequency in the lab strains and wild population of *Drosophila affinis*. Mean proportion is the average proportion of XO male frequency across individual fathers, SE refers to standard error calculated across those fathers. Overall proportion is total XO sons divided by males screened. SEP refers to the standard error of the proportion.

| Group | Broods screened | Males screened | XO sons | Mean prop. XO (SE) | Overall prop. XO (SEP) |
| --- | --- | --- | --- | --- | --- |
| Lab $X^{SR}Y$ males | 51 | 56 | 42 | 0.775 (0.075) | 0.750 (0.058) |
| Lab $X^{ST}Y$ males | 69 | 313 | 0 | 0.0 (NA) | 0.0 (NA) |
| Wild males | NA | 165 | 3 | NA | 0.018 (0.010) |

We performed a similar crossing scheme to measure the rate of nondisjunction in standard *darkeye* males (but crosses with virgin *genome* females to distinguish Y chromosome from the X nondisjunction based on eye color). In total 69 broods produced offspring, and approximately 46% (1459 out of 3171) of the offspring were male. We tested 313 total males, and not a single XO male was detected (Table S.2). Using Hanley’s method to estimate a 5% upper confidence limit for a proportion with 0 positives and 313 total tested, we get a range from 0 to 0.95% nondisjunction in standard *darkeye* males (Hanley and Lippman-Hand, 1983). For the 313 males, we first took all sons from 11 clutches, and none of the 77 males were XO males. So we then randomly chose a total of 236 males from the rest of 58 broods, again not a single XO male was detected. Finally, the rate of nondisjunction in sex-ratio males (75%) is significantly higher than that in standard *darkeye* males (0%) (Fisher’s exact test, p < 2.2e-16).

#### 3.1.2 The frequency of XO males in a wild population

To explore the frequency of XO male in the wild population, we used males collected at the Rees’ Fruit Farm near Topeka, KS in August 2021. Among the 165 male samples, 3 males were classified as XO males (1.8%, (Table S.3). If we assume strict mutation-selection balance in standard males, with a nondisjunction rate of 0.1%, and strong selection against the O (*s*_*O*_ = 0.25) (Voelker and Kojima, 1972), the expected frequency of XO males in the wild should be about 0.4% (*u*/*s*_*O*_ = 0.001/0.25). Thus the observed frequency in the wild is more than 4 times the frequency expected under mutation-selection balance (*χ* ^2^ = 7.555, df = 1, p = 0.006). However, the expected XO frequency in wild flies is difficult to estimate due to various uncertain but crucial factors including the cost of O in XO males.

### 3.2 A model of O resistance to sex-ratio meiotic drive

#### 3.2.1 Empirical estimates of nondisjunction rates in X^SR^ and X^ST^ (standard X chromosome, without driver) males

The results from lab experiments to estimate nondisjunction rate suggest that it is rare, but must be nonzero in X^ST^ males (*µ* ≈ 0.001) given what has been measured for other *Drosophila* species (Bridges, 1913; Tokunaga, 1970). The nondisjunction rate in X^SR^ males is much higher (*λ* ∗ *µ* ≈ 0.75) - likely due to the drive mechanism itself. This gives us an estimate of *λ* ≈ 750. In other words, nondisjunction rates are approximately 750 times higher in X^SR^ males compared to X^ST^ males.

Moving forward, we will discuss our model using the full parameterization (see methods and appendix) and also a simplified, idealized model where we assume SR costs in female are homozygous lethal (*h* = 0 and *s*_*X SR*_ = 1), drive strength in X^SR^O males is fully reversed (all sons produced, *d*_*O*_ = −0.5), and the X^SR^ chromosome has no cost in males (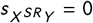 and 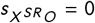). This simplified model both allows for analytical solutions (which we have not been able to obtain with the full model) and presents results that are easier to intuit.

#### 3.2.2 The fate of the O and X^SR^ chromosome in isolation

We begin by considering the fate of the O and X^SR^ chromosome individually. This is straightforward and has been considered elsewhere (Hartl, 1970; Crow, 1991; Hall, 2004), but we rederive here with the nuances of our specific model. First, in the absence of sex-ratio meiotic drive, if the O carries any cost, it will segregate at mutation-selection balance (Hartl et al., 1997) where mutation here refers to the rate of nondisjunction (*µ*) and selection (*s*_*O*_) is the cost of being X^ST^O relative to X^ST^Y. Therefore, in the absence of drive, the O segregates at a frequency of 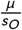. Assuming nondisjunction occurs at rates around 0.001 (one per 1000 meioses) and selection against XO is strong (*s*_*O*_ ≈ 0.25 (Voelker and Kojima, 1972)), the O should segregate at a frequency of about 0.4% in the absence of meiotic drive.

Next, we consider the frequency of X^SR^ chromosome (X^SR^) in the absence of the O (or any Y-linked resistance). The full solution is in the supplemental text (Equations A.13 and A.14). In our simplified model, the internal equilibrium (denoted by the caret) occurs when:

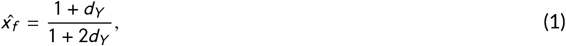

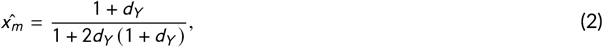

Equation 1 provides an easy to intuit picture of how allele frequencies change with the strength of drive (in females). Without drive (*d*_*Y*_ = 0), the equilibrium frequency of the standard X equals one (X^SR^ equals zero) since there is no drive. When drive is full (*d*_*Y*_ = 0.5), the frequency of the standard X equals 0.75 (X^SR^ equals 0.25), with X^SR^ frequency increasing in a concave down pattern.

The frequency of the O at mutation-selection balance provides a baseline frequency when asking whether a new X^SR^ chromosome can invade when the O is segregating (Equation 2). The equilibrium frequency of X^ST^ (and therefore X^SR^) provides a baseline for determining the effect of the O (and rates of nondisjunction) on the equilibrium frequency of X^SR^.

#### 3.2.3 Invasion of X^SR^ when O is segregating

Our first goal is to determine the parameters allowing for the invasion of the X^SR^ chromosome when the O is segregating at mutation-selection balance. We get a reasonable estimate of the critical strength of drive for invasion of the X^SR^ chromosome which is when X^SR^’ > X^SR^ in males - when the frequency of X^SR^ in the next generation is greater than its frequency in the current generation. The focus on males simplifies the solution, but also seems reasonable since fitness consequences in females only matter if they are dominant, when X^SR^ is rare. We additionally assume that we start with a single X^SR^ allele, meaning that the frequency of X^SR^ is 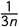 in each sex and the O is at mutation-selection balance. The full solution (Equation 3) is that the X^SR^ chromosome can invade when:

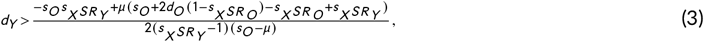

Therefore, X^SR^ invasion is mostly dependent on the cost of X^SR^ in males (which balances the benefit to the chromosome of driving). Using the idealized scenario, this simplifies to (Equation 4):

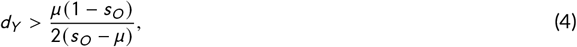

With very low cost associated with being XO (*s*_*O*_), drive must be very strong (*d*_*Y*_ > 0.4) for invasion because an O with low cost will segregate at higher mutation-selection balance frequencies. However, the strength of drive necessary for invasion drops quickly and asymptotically approaches zero with increasing cost of being XO.

In the absence of the O, the driver can initially invade as long as 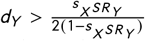. Figure 2 shows the X^SR^ invasion criteria in the idealized scenario and when the cost of drive in males 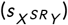 is 0, 0.2 and 0.4. In the absence of the O, invasion requires the strength of drive (*d*_*Y*_) to be greater than 0, 0.125 and 0.333 for those three costs in males, respectively. Figure 2 also makes it clear that the O only prevents invasion of X^SR^ when its frequency in the standing genetic variation (*µ*/*s*_*O*_) is relatively high. This requires nondisjunction to be near 1% in normal males and selection against the O to be relatively weak (*s*_*O*_ < 0.25).

**FIGURE 2.**
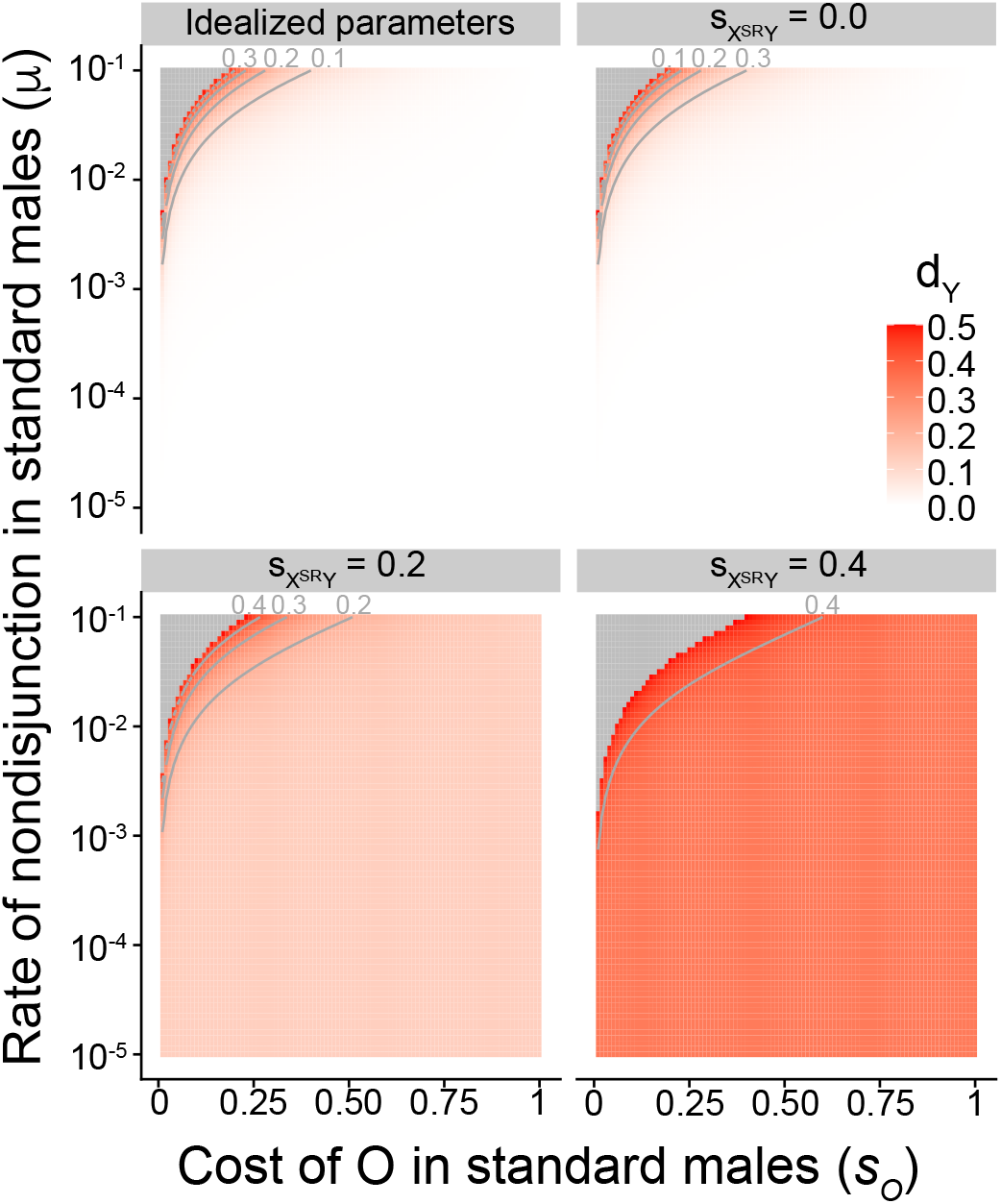
Invasion of X^SR^ with the O segregating at mutation-selection balance. The strength drive (*d*_*Y*_, indicated by the shade of red) required for X^SR^ invasion when starting from a single individual and O at mutation-selection balance. See text for idealized parameters. Other plots assume 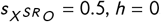, and *d*_*O*_ = −0.5. Contours indicate increments of 0.1.

The invasion of the O is of little interest since we assume it is segregating at mutation-selection balance and therefore has already “invaded”, albeit at low stable equilibrium frequency.

#### 3.2.4 Equilibrium frequencies of X^SR^ and O

After invasion, X^SR^ and O may stably oscillate or reach a stable equilibrium. We were unable to solve the system of equations (see Appendix Equations (A.1)-(A.3)) with all parameters, but in the idealized scenario, there are four equilibria. The first occurs when the standard X and O are both fixed. The second occurs when the standard X is fixed and the O is at mutation-selection balance. The third is several lines of formula but results in internal equilibria for both X^SR^ and O, and the fourth does not lead to a reasonable (between zero and one) frequency of the O with our parameters. Due to the complexity of the result, we do not spell out the internal equilibrium here (see Appendix Equations A.10-A.12), but the values are presented in Figure 3 for the O, the frequency of X^SR^ in females, and the relative reduction of X^SR^ frequency with the O segregating compared to a fixed Y. Although we solve for the frequency of the standard X (X^ST^) and the Y, these plots show the frequencies of the X^SR^ chromosome and the O since they are the variables of interest. Simulation results confirmed these equilibrium values (Table S.4).

**FIGURE 3.**
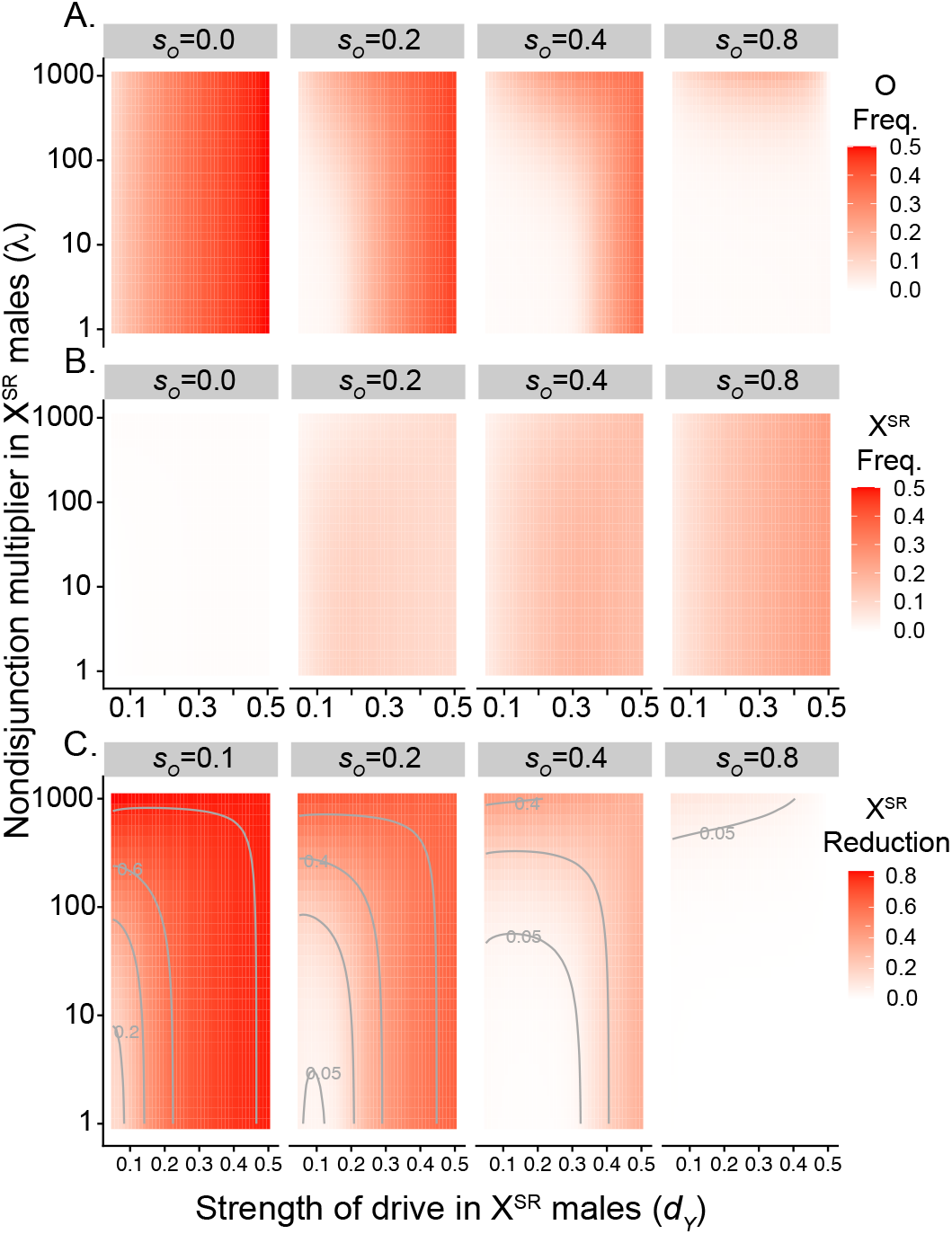
Equilibrium frequencies in the idealized scenarios. A) Equilibrium frequency of the O, B) equilibrium frequency of X^SR^ in females, and C) the relative reduction in X^SR^ frequency in females when O is segregating compared to a fixed Y. The rate of nondisjunction in XY males is assumed to be *µ* = 0.001.

Figure 3 demonstrates that cost of the O in XO males (*s*_*O*_), strength of drive (*d*_*Y*_), and the nondisjunction multiplier (*λ*) all influence both O and X^SR^ frequency. When the cost of the O is low, and drive strength and nondisjunction are both high (3C, leftmost panel), the frequency of the driver can be reduced by nearly 80%.

#### 3.2.5 Stability versus oscillations

In the idealized case, the equilibria are stable. However, when we relax the assumptions of the idealized case, we see more complex behavior. To explore parameter space, we conducted deterministic simulations to ask whether the system reached a stable equilibrium or showed oscillated indefinitely. We ran these simulations for 20,000 generations which was enough time for any parameter set we examined to either reach a stable equilibrium or settle into stable oscillations. We defined stability as having a variance in allele frequency of less than 0.0001 during the final 2000 generations of the simulation. Figure 4A demonstrates how the homozygous cost SR in females leads to different types of stability. With low cost, (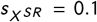), both X^SR^ and O oscillate wildly (but stably): X^SR^ ranging from close to zero to more than 0.5 and O ranging from close to zero to close to one. With intermediate cost 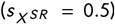, oscillations dampen and but never reach a single stable value. Finally, with high cost 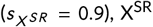 and O reach stable values quickly with little to no oscillation.

**FIGURE 4.**
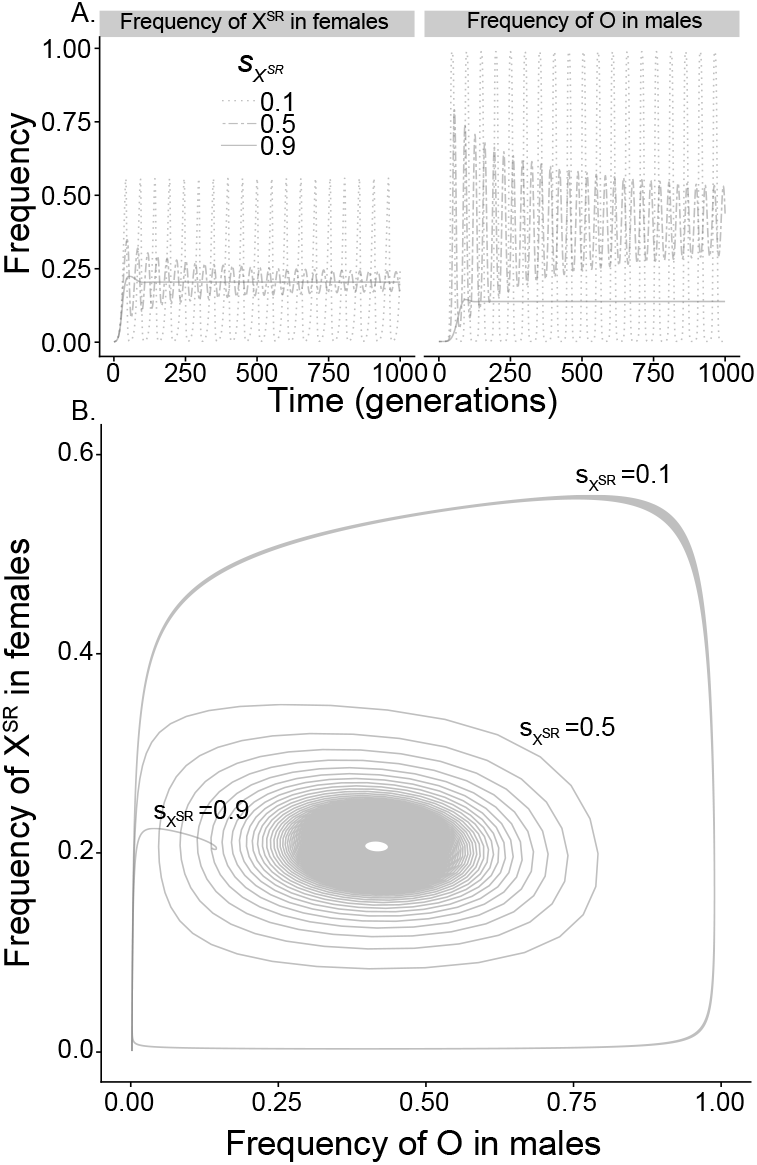
Lower cost of X^SR^ in females leads to oscillating allele frequencies. A) Representative trajectories for X^SR^ and O with homozygous cost of X^SR^ ranging from 0.1 to 0.9 (1000 generations displayed), B) The same data with ferquencies of X^SR^ in females and O in males plotted on the Y and X axis, respectively (10,000 generations displayed). Other parameters for these simulations: 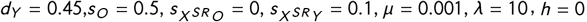, and *d*_*O*_ = −0.5. Simulations begin with the O at mutation selection balance (*µ*/*s*_*O*_) and the frequency of X^SR^ = 0.001 in both sexes.

The same results are plotted in Figure 4B, where now the frequency of X^SR^ in females is plotted against the O. The stable oscillations at low cost in females, dampening oscillations with intermediate cost, and stable equilibrium at high cost are clear in these plots. Note specifically that with intermediate cost 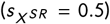, oscillations dampen, but then settle into a stable oscillation. Figure S.1 shows more generally where behavior changes from oscillations to stable equilibrium with a wide range of parameters. Overall, homozygous cost in female 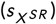 has the most influence on this behavior with most parameter space shifting from oscillations to stable equilibrium when costs are moderate 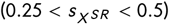. All simulation results for the general case are presented in Table S.5.

#### 3.2.6 Sex chromosome turnover

A critical question is whether or not the combination of driver and O would ever lead to the complete loss of the Y chromosome. When the equilibrium is stable, this seems unlikely - the O rarely reaches equilibrium frequencies above 0.5. This is because once the driver is at very low frequency (or lost), the O has no further benefit and therefore cannot become fixed. However, oscillations in allele frequency may lead to Y frequencies so low that it would in fact be lost in any finite population. This has significant implications. First, it would lead to the extinction of the Y chromosome. This would, at least temporarily, lead to the transition from XY to XO sex chromosome system. Subsequently a B chromosome or autosome, that improves male fitness and segregates against the X, might establish a new Y, leading to a complete turnover of the Y chromosome. Second, as soon as the Y is lost, X^SR^ has no selective advantage and is quickly purged from the population. So fixation of the O means loss of both Y and X^SR^.

To determine parameters that would lead to the extinction of the Y, we stopped simulations if the frequency of the Y was less than 1/*N*_*m*_ where *N*_*m*_ is the effective population size of males which we assume to be 10,000. However, this never led to extinction of the Y - though O oscillations ranged from very low frequency to very high frequency (>0.99), we did not see frequencies that would be associated with fixation of the O in a moderately sized population (*N*_*m*_ = 10000 or even much smaller). In fact, it wasn’t until we increased the cost of being XO to greater than 0.9 and decreased the cost of being homozygous 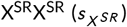 that we ever saw the Y drop to less than 1/*N*_*m*_. These parameters seem unlikely given what we know about the *D. affinis* system (personal observation). Reports on the cost of the XO are mixed. Volker et al. (Voelker and Kojima, 1972) found that selection coefficients against the O to be high (25-36%), but Unckless et al. (2015) observed no cost in terms of mating components. The discrepancy could be due to the fact that different phenotypes were measured (selection in a cage over time vs. specific mating success) or because the Y and O were on different genetic backgrounds in the two studies. In the lab, homozygous X^SR^X^SR^ females have show an approximately 75% reduction in offspring production compared to X^SR^X^ST^ females. Frequencies when the O reaches a stable value are shown in Figure S.2 indicating that under no circumstances did the O reach anything near fixation. The complete set of simulation results are in Table S.6.

## 4 DISCUSSION

Organisms are impressively complex systems that have evolved over millions of years to function in a delicate homeostatic balance. However, the genomes of these organisms are also fraught with selfish genetic elements that poke and prod at this balance to improve their own fitness without regard to the fitness of the host. Therefore, it should not be surprising that sometimes the selfish genetic elements push and prod too far and the result is to their own detriment. For example, some selfish genetic elements alter sex ratios to such extremes that entire populations go extinct (Hamilton, 1967; Jiggins et al., 2002; Pinzone and Dyer, 2013; Unckless and Clark, 2015). Here we describe a new example of a selfish genetic element getting in its own way: sex-ratio meiotic drive is associated with dramatically higher rates of nondisjunction of the Y chromosome in *D. affinis* and males lacking a Y chromosome are not only fertile, but also resistant to drive (Figure 1). This type of resistance arising from the natural drive machinery is reminiscent of what occurred in early models of homing gene drives (Burt, 2003; Deredec et al., 2008; Hammond et al., 2016; Champer et al., 2016; Unckless et al., 2017), and understanding the mechanisms and evolution of resistance in natural systems is crucial for engineering effective gene drive systems.

Sex determination in *Drosophila* is independent of the Y chromosome, but the Y is required for fertility in most species. Though sex-ratio meiotic drive is associated with nondisjunction of the Y chromosome in other species (Henahan and Cobbs, 1983; James and Jaenike, 1990; Dyer, 2012), XO males in those species are sterile, so the O is an evolutionary dead end. In *D. affinis*, the fact that XO males are fertile and X^SR^O males produce exclusively sons, means that the O can be selected upon and potentially even spread through populations resulting in the loss of the Y chromosome. Nondisjunction in SR males of both *D. pseudoobuscura* and *D. neotestacea* was complete - all sons of SR males were sterile XO (Henahan and Cobbs, 1983; James and Jaenike, 1990; Dyer, 2012). Y chromosome nondisjunction in *D. affinis* SR males, however, is not quite complete (about 75%). Although we did not detect nondisjunction in the sons of standard (XY) males (0 out of 313), the rate in SR (X^SR^Y) males was about 75%. If we assume Y chromosome nondisjunction rates are between 0.05 and 0.1% in standard males as in (Bridges, 1913; Tokunaga, 1970), this corresponds to a 750 times higher rate of nondisjunction in SR males. Note that in addition to our method of PCR detection of XO males, we might have performed chromosome squashes to screen for the presence/absense of the Y. Unfortunately, the DNA extractions for these males are destructive, so we cannot easily test both DNA and karyotype, and karyotyping all individuals would be prohibitive.

The actual molecular causes of drive are only partially understood in these *Drosophila* SR systems (Pieper et al., 2018), and it may be that these differences reflect fundamentally different mechanisms of drive. *D. affinis* Y chromosomes also show variation in susceptibility to drive - with some complete resistance (Unckless et al., 2015). In this extreme case, we know that most sons of X^SR^Y^resistant^ males do inherit the resistant Y (or else that Y would quickly be lost in the population and in stocks). In contrast, Y-linked resistance to SR meiotic drive is lacking in both *D. pseudoobscura* and *D. neotestacea* (Lindholm et al., 2016). It would be interesting to determine how the rate of male production in *D. affinis* correlates with the rate of Y chromosome nondisjunction for these different Y chromosomes, and in turn, the implications of nondisjunction on the spread of partially resistant Y chromosomes.

Our modeling approach demonstrates that the O likely impacts SR meiotic drive dynamics in three ways. First, an O segregating at mutation-selection balance can prevent X^SR^ from invading a population (Figure 2). The conditions for this are relatively restrictive unless drive is costly in males (which would generally restrict the drive’s ability to invade). However, the parameters that would prevent invasion of X^SR^ even in the idealized model (top left Figure 2) are not completely out of the question. While Voelker and Kojima (1972) inferred strong selection against the O (*s*_*O*_ > 0.25), Unckless et al. (2015) found no evidence for fertility differences between XY and XO males (in the absence of meiotic drive), so the fitness of the O may be largely dependent on autosomal background and average *s*_*O*_ may actually be quite low. Furthermore the rate of nondisjunction likely depends on environmental, age, and genetic factors. In *D. melanogaster*, low temperature can lead to nondisjunction rates as high as 7% Tokunaga (1970). *D. affinis* ranges from the USA/Canada border south into Florida and Texas (Miller, 1958), so it experiences broad seasonal and clinal temperature range which could lead to clinal O frequencies, and potentially as a consequence, clinal X^SR^ frequencies. Other species show such clinal patterns in meiotic driver frequencies, but they lack the O and the most common explanation is correlation with other life history factors (*e*.*g*. pleiotropic selection on the X chromosome or polyandry) (Pinzone and Dyer, 2013; Price et al., 2014; Stewart et al., 2019).

The second impact of the O is that it can lead to stable oscillations, particularly when the cost of SR in homozygous females is low (Figures 4 and S.1). This result, however, is not new and was reported by Hall (2004) for Y chromosomes showing complete resistance (but not reversal) of drive. Finally, the O can significantly reduce the equilibrium frequency of X^SR^ (Figure 3). This effect can be striking, particularly if the drive strength is high (*d*_*Y*_ > 0.4) and costs in XO males are low (*s*_*O*_ < 0.25). Our anecdotal estimates for *D. affinis* are that costs of X^SR^ in females are high and recessive 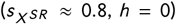, drive is strong (*d*_*Y*_ ≈ 0.486, costs of X^SR^ in males is low 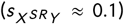, costs of X^SR^ and O in males is high 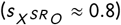 and the nondisjunction multiplier (*λ*) is about 750. Our deterministic simulation with these parameters leads to an equilibrium driver frequency in females of 0.367 and an O frequency of 0.062. The O frequency is somewhat higher than what we observed in the wild population (0.018, Table 2). However, this X^SR^ frequency is much higher than our observed driver frequency in wild populations (0.02 to 0.10, personal observation), so other factors including Y-linked resistance must be at play.

One potential consequence of both X^SR^ and O segregating in a population is the loss of the Y chromosome, or the loss of X^SR^. Our simulation results suggest that this is unlikely unless parameters are much more extreme than what we observe for *D. affinis* or other systems. For instance, our simulations rarely reached above 99% which would mean that even population as small as Ne=100 would not experience the loss of the driver or the Y. However, the O could lead to sex-chromosome turnover indirectly. We speculate that an X-autosome fusion would also lead to a neo-Y chromosome segregating against the autosomal homolog of the neo-X. This neo-Y chromosome a) is likely not sensitive to drive or even results in all sons as in X^SR^O males (since it wasn’t targeted as an autosome), and b) could then accumulate male-beneficial mutations. Thus the neo-Y could complete the displacement of the ancestral Y because, unlike the O, it can accumulate male beneficial mutations. This scenario of meiotic drive and the O leading to sex chromosome turnover is similar to that proposed by van Doorn and Kirkpatrick (2007) where sexually antagonistic selection drives sex chromosome turnover. Interestingly, the Y chromosome shows size variation in *D. affinis* and several other members of the Obscura group of *Drosophila* (Dobzhansky, 1935; Miller and Stone, 1962). It would be interesting to investigate whether these Y chromosomes have gene translocation from other autosomes, which may hint at the initial stages of sex chromosome turnover. It seems unlikely that the O would lead to the loss of the X^SR^, though it may prevent establishment in the first place (in specific conditions).

Our results are similar to what Bravo Nũnez *et al*. (2020) describe in fission yeast. The fission yeast *wtf* drive systems usually involve both toxin and antidote (encoded by two isoforms of the same gene). The toxin is secreted and kills any spores except those that produce the non-secreted antidote. *Schizosaccharomyces pombe* isolates are so rife with autosomal meiotic drivers (*wtfs*) that heterozygotes often have two different drivers, each poised to kill spores bearing the homolog. Therefore, with normal chromosome segregation all products of meiosis perish. If there is nondisjunction of the autosomes containing the driver, half of the spores will lack that autosome and therefore be inviable. The other half, however, inherit both drivers and both antidotes and therefore survive. Thus, the authors demonstrated that *wtf* drivers could generate atypical diploid spores (through nondisjunction) that were advantageous in the meiotic drive context. Both genetic work and population genetic modeling suggest that mutations increasing the rates of nondisjunction can be favored in this system. So in both yeast and flies, nondisjunction may provide an escape from selfish genetic elements. Note however, that we suspect a poison/target model (as in *sd* in *D. melanogaster* (Larracuente et al., 2010)) in *D. affinis*, not the poison/antidote system described for *S. pombe*, which is why *X* ^*SR*^ *O* males avoid the toxin - they have no target.

In the grand scheme, *Drosophila* is somewhat unfertile ground for this type of dynamics since the Y is required for fertility in all but a very few species (Voelker and Kojima, 1971). Other taxa have much more diversity in terms of sex chromosome identity and some likely tolerate a lack of Y chromosome much better than most *Drosophila* species. For instance, the Y is often required for fertility in the vast majority of mammals, except the mole vole (*Ellobius lutescens*), *Tokudaia* genus rodents, and the creeping vole (*Microtus Oregoni*, with XO female sex-determining system) with natural XO male sex-determining system (reviewed by (Saunders and Veyrunes, 2021)). In addition, some groups such as Coleoptera and Diptera, have multiple independent losses of the Y chromosome (Blackmon and Demuth, 2014), and certain Lepidoptera insects lost the W chromosome and became a ZO sex-determining system (Traut et al., 2008). It would be interesting therefore to investigate whether the incidence of drive is associated with presence/absence of a Y chromosome among different taxa. When does the evolutionary advantage of a male-specific chromosome for male-specific functions such as male fertility not justify the added liability associated with being a target for genetic conflict?

## Acknowledgements

We thank Elizabeth R. Everman and Paris Veltsos for the help with wild male *D. affinis* collection, Hunter L. Duke for help with some of the fly counting. The title came out of discussion with Sarah Zanders. This work was funded by NSF CAREER grant (2047052) to RLU.

## Author Contributions

RLU and WJM designed the research. WJM did the fieldwork. WJM, EMK, KBP, and MMS conducted the related crosses. WJM, EMK and KBP conducted DNA extractions and PCR to screen for the frequency of XO males. RLM performed the modeling work, and contributed reagents and funding. RLU and WJM wrote the manuscript. All authors approved the final version of the manuscript.

## A APPENDIX

### A.1 A sex-ratio meiotic drive model with nondisjunction

We begin with a model similar to Hall (2004). Parameters and variables are describe in Table 1. The system is described by the following recursions and mean fitness values.

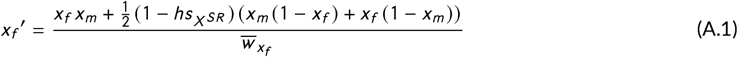

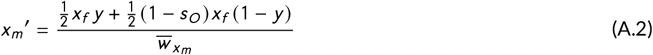

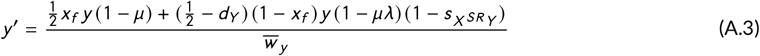

where

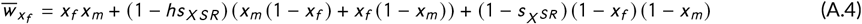

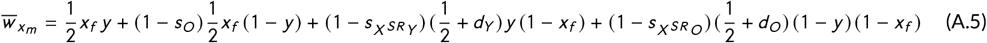

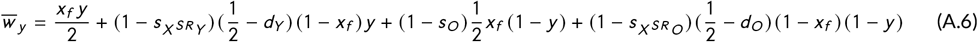

We assume that fitness costs are experienced before nondisjunction. There are several differences from the (Hall, 2004) model. Most of these involve fitness costs in males. However, we also allow for a background rate of nondisjunction (*µ*) of the Y in standard males, then a multiplier of that nondisjunction rate (*λ*) in SR males. The multiplier, *λ*, can range from 1 (the same rate as in standard males) to 1/*µ* where the sons of SR males are always XO.

We were not able to solve A.1-A.3 analytically, so we derived an idealized model where we assume fitness costs in females are recessive lethal (*h* = 0 and 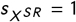), all offspring of X^SR^O males are sons 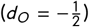, and there are no costs associated with X^SR^Y or X^SR^O males. These assumptions simplify the recursions in A.1-A.3 and the associated mean fitness values to:

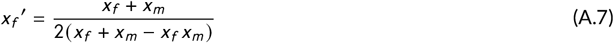

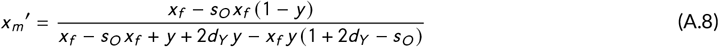

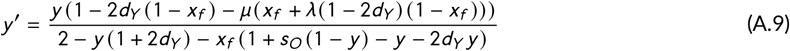

Equations A.7-A.9 include normalization to mean fitness.

### A.2 Equilibrium frequencies

We solved the system of equations in A.7-A.9 for equilibrium (i.e. when 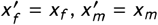 and *y* ′ = *y*). There are four solutions. The first is when the standard X is fixed (*x*_*f*_ = *x*_*m*_ = 1) and the O is fixed (*y* = 0). The second is when the standard X is fixed and the O segregates at mutation-selection balance. The third and forth are messy equations that do little in terms of informing intuition, and only the third leads to equilibrium values between 0 and 1, so we focus on that here. Those values are:

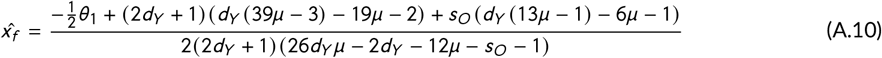

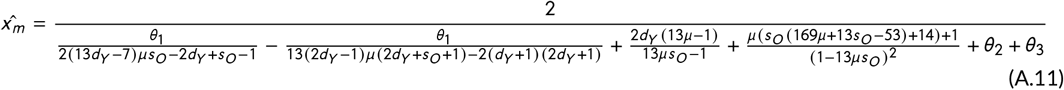

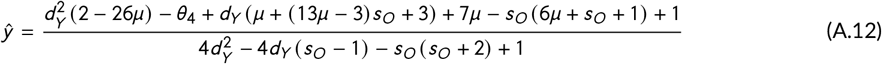

where

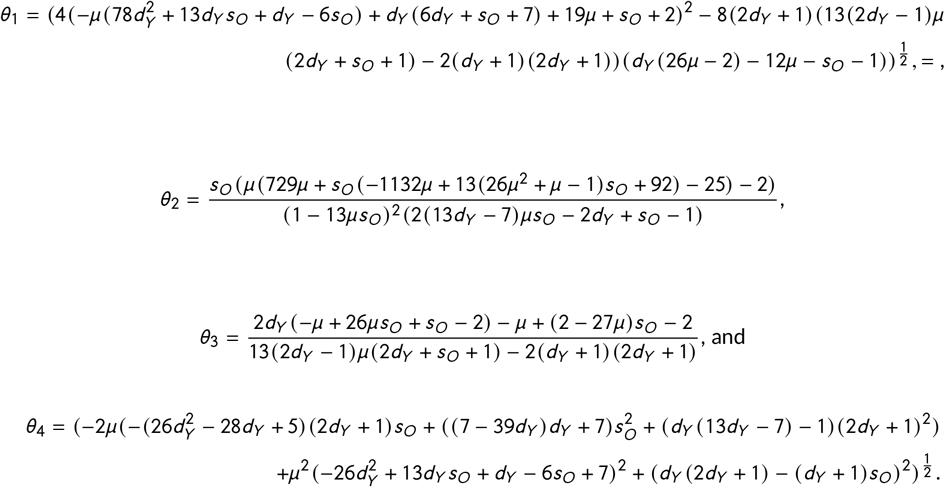

#### A.2.1 Equilibrium frequencies of X^SR^ when O is absent

For the full model, we found the equilibrium frequencies of the X when the Y is fixed (O is not segregating). There are three solutions, two of which are fixation of the standard or SR X chromosome. The third solution is:

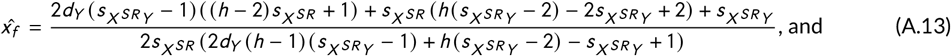

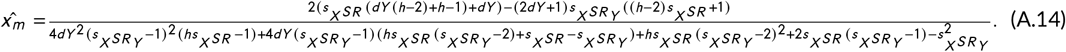

The equilibrium conditions for the idealized model are presented in the main text Equations 1 and 2.

## Supplement

### Tables

**TABLE S. 1** PCR Primers used in this study to amplify a Y-linked locus and the *COI* locus. All sequences are presented in the 5’ to 3’ direction.

**TABLE S. 2** Brood information and nondisjunction rates for individual males from the lab. Data includes a brood identifier, father’s genotype, mother’s genotype, the number of total females, number of males, number of tested males, proportion of females, number of XO males and proportion of males that are XO. Each male was mated to 1-4 females homozygous for the *darkeye* mutation.

**TABLE S. 3** PCR results for wild-collected *D. affinis* males. All males were PCR screened for a Y-linked locus and *RCC1* locus. The *RCC1* locus PCR was to ensure DNA quality, and those samples with repeated absence of Y-linked locus were considered as XO males.

**TABLE S. 4** Simulation results for idealized parameters. “t” refers to the generation at which we stopped the simulation, “Stability” is 1 if the final 2000 generations of the simulation had a variance of less than 0.0001, “Invade” refers to whether the frequency rose above 0.01, and if not, the highest frequency observed.

**TABLE S. 5** Simulation results for general parameters. “t” refers to the generation at which we stopped the simulation, “Stability” is 1 if the final 2000 generations of the simulation had a variance of less than 0.0001, “Invade” refers to whether the frequency rose above 0.01, and if not, the highest frequency observed.

**TABLE S. 6** Simulation results for determining whether the Y (or X^SR^) go extinct. “t” refers to the generation at which we stopped the simulation, “Stability” is 1 if the final 2000 generations of the simulation had a variance of less than 0.0001, “Invade” refers to whether the frequency rose above 0.01, and if not, the highest frequency observed.

### Figures

**FIGURE S.1.**
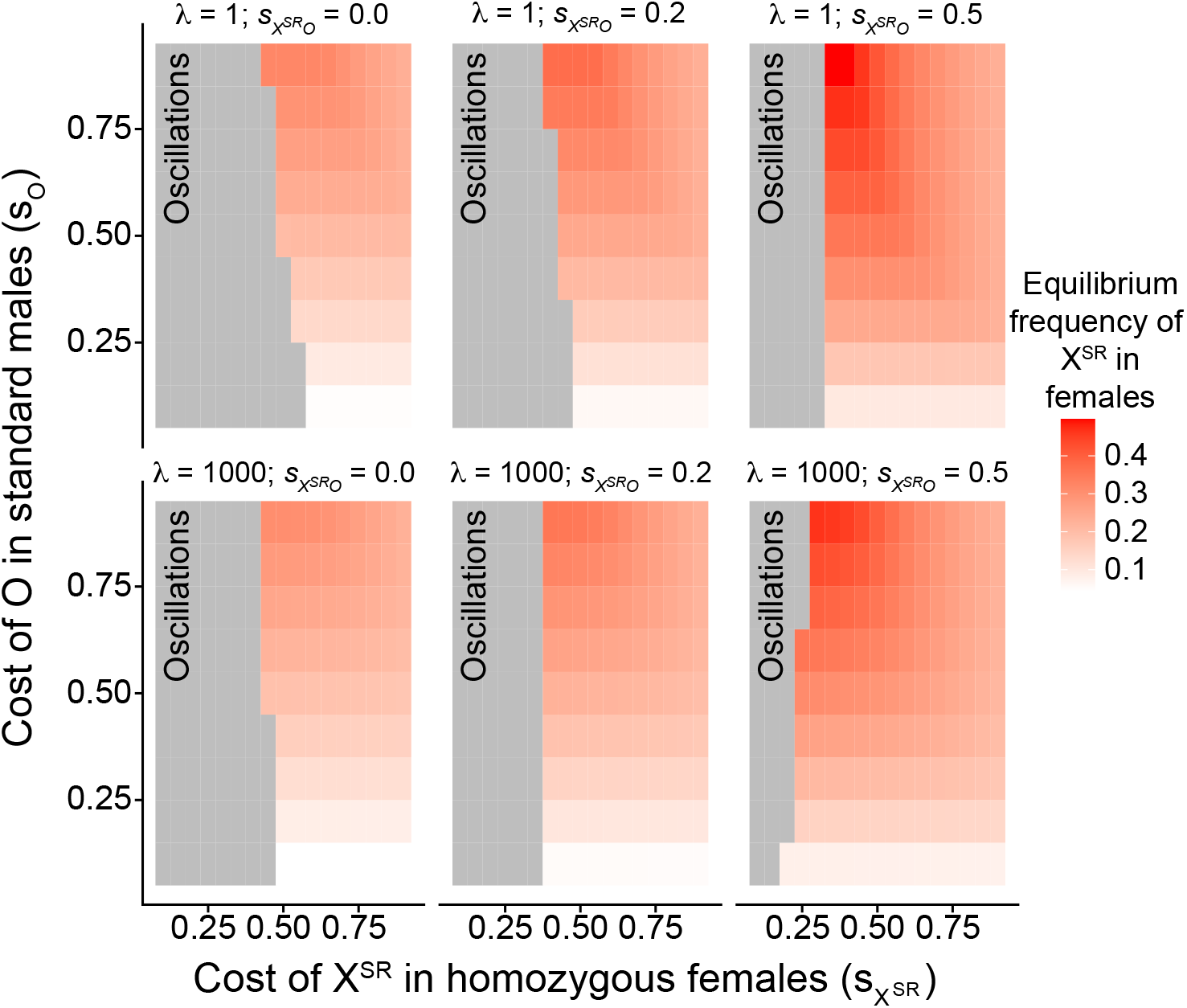
Equilibrium frequency and stability with low (*λ* = 1) and high (*λ* = 1000) nondisjunction multipliers in X^SR^Y males, no, medium or high costs (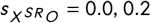 or 0.5) in X^SR^O males, and a range of costs in XO males (*s*_*O*_) and homozygous SR females 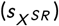. In all cases, costs in females are recessive (*h* = 0), the baseline nondisjunction rate (*µ*) is 0.001, the strength of drive (*d*_*Y*_) is 0.45, cost in SR males 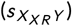 is 0.1, and the strength of drive in X^SR^O males is (*d*_*O*_) -0.5 (all sons).

**FIGURE S.2.**
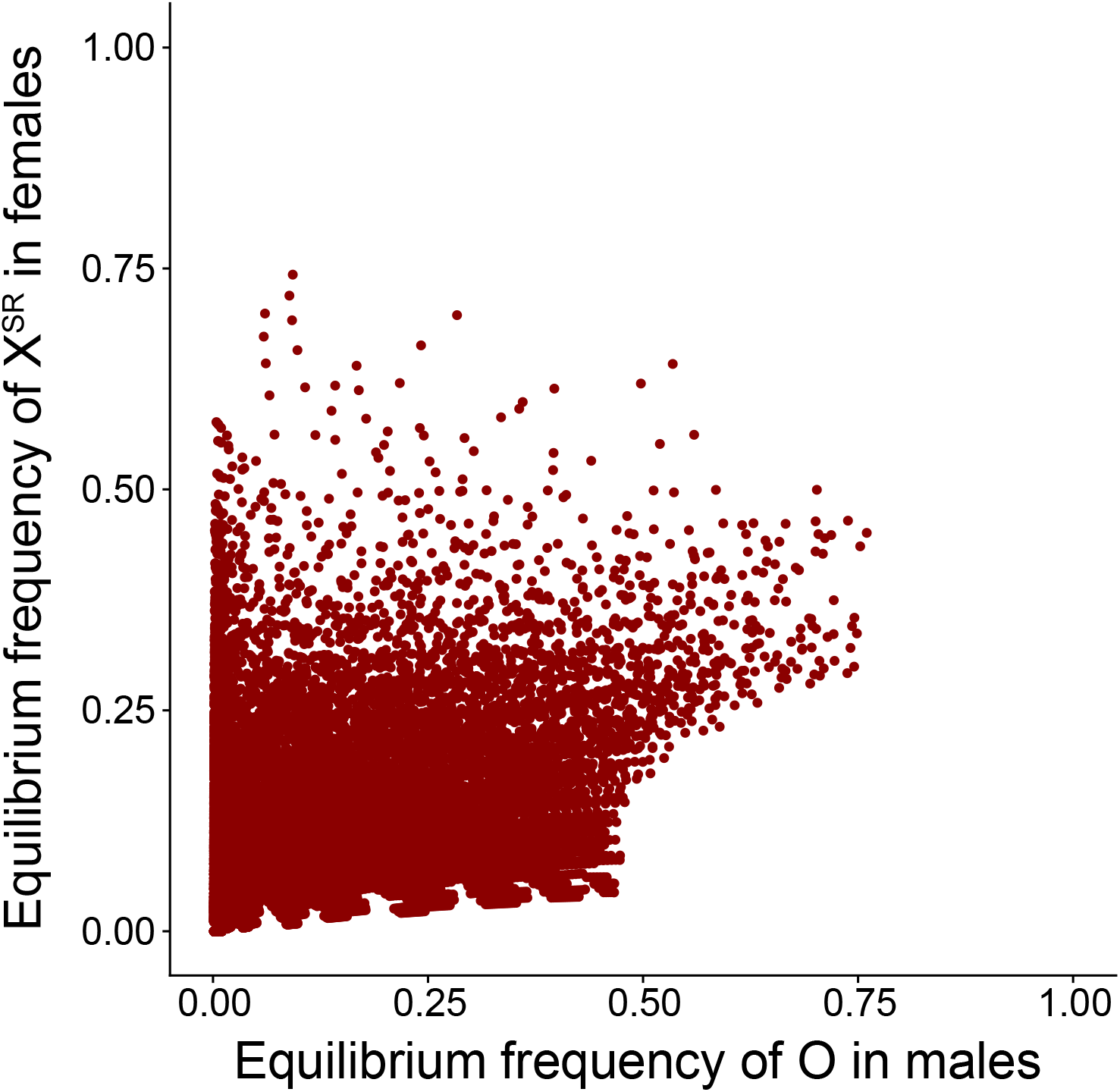
Neither the O nor X^SR^ chromosome are driven extinct in this system. Equilibrium frequency of X^SR^ in females and the O in males as determined via simulation with the following fixed parameters: costs in females are recessive (*h* = 0), the baseline nondisjunction rate (*µ*) is 0.001, and the strength of drive in X^SR^O males is (*d*_*O*_) -0.5 (all sons). Variable parameters are the strength of drive (*d*_*Y*_ ranging from 0.05 to 0.45 in increments of 0.1), cost in SR males (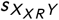 ranging from 0 to 0.5 in increments of 0.1), nondisjunction multiplier (*λ* as 1 or 1000), costs in X^SR^O males (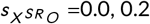 or 0.5), costs in XO males (*s*_*O*_ ranging from 0.1 to 0.9 in increments of 0.1) and costs in homozygous SR females (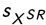 ranging from 0.1 to 0.9 in increments of 0.05).

